# Variable impact of geochemical gradients on the functional potential of bacteria, archaea, and phages from the permanently stratified Lac Pavin

**DOI:** 10.1101/2022.07.18.500538

**Authors:** Alexander L. Jaffe, Corinne Bardot, Anne-Hélène Le Jeune, Jett Liu, Jonathan Colombet, Fanny Perrière, Hermine Billard, Cindy J. Castelle, Anne-Catherine Lehours, Jillian F. Banfield

## Abstract

Permanently stratified lakes contain diverse microbial communities that vary with depth, and so serve as useful models for studying the relationships between microbial community structure and geochemistry. Recent work has shown that these lakes can also harbor numerous bacteria and archaea from novel lineages, including those from the Candidate Phyla Radiation (CPR). However, the extent to which geochemical stratification differentially impacts carbon metabolism and overall genetic potential in CPR bacteria compared to other organisms is not well defined. Here, we determine the distribution of microbial lineages along an oxygen gradient in Lac Pavin, a deep, stratified lake in central France, and examine the influence of this gradient on their metabolism. Genome-based analyses revealed an enrichment of distinct C1 and CO_2_ fixation pathways in the oxic lake interface and anoxic zone/sediments, suggesting that oxygen likely plays a role in structuring metabolic strategies in non-CPR bacteria and archaea. Notably, we find that oxidation of methane and its byproducts is largely spatially separated from methane production, which is mediated by diverse communities of sediment methanogens that vary on the centimeter scale. In contrast, we detected evidence for RuBisCO throughout the water column and sediments, including form II/III and form III-related enzymes encoded by CPR bacteria in the water column and DPANN archaea in the sediments. Overall, CPR bacteria and phages did not show strong signals of gene content differentiation by depth, despite the fact that distinct species groups populate different lake and sediment compartments. Thus, environmental gradients in Lac Pavin probably select for the capacities of CPR bacteria and phages to a lesser extent than other bacteria and archaea, possibly because selection on the former groups is indirect and depends on host characteristics.

## INTRODUCTION

Meromictic lakes are permanently stratified aquatic ecosystems with a global distribution [1]. Despite variation in size and morphology, these lakes generally have several common physical features, including the presence of two major layers - a shallower, oxygenated layer called the mixolimnion, and a deeper, generally anoxic zone called the monimolimnion [2]. These layers are separated by an oxycline of variable steepness that occurs in the water column, forming an ‘interface’ zone. A unique feature of meromictic lakes, compared to other similar bodies of water, is that mixing events between the two layers are rare, with seasonal mixing observed primarily within the oxygenated upper layer [2]. Based primarily on marker gene analyses, these compartments are known to host distinct and diverse microbial communities that co-vary with gradients in oxygen, light, and sulfur [3,4], and thus can serve as model systems for probing the relationships between community structure, metabolism, and geochemistry.

Archaea and bacteria with the capacity to utilize one-carbon (C1) molecules as carbon and energy sources are widespread across the planet and are important players in many biogeochemical processes. As such, they occupy an array of diverse anaerobic and aerobic niches and perform unique metabolisms that enable the utilization of carbon dioxide (CO_2_), carbon monoxide (CO), formate (HCOOH), methanol (CH_3_OH), and methane (CH_4_) for growth, including in meromictic lakes. However, the spatial distribution and phylogenetic affiliation of pathways associated with carbon compound transformation over the water column and sediments of these lakes is still emerging. Previous work has also shown that these ecosystems can host organisms from divergent lineages without representatives in pure culture, including the Candidate Phyla Radiation (CPR) bacteria and a novel sibling lineage of the Cyanobacteria (Ca. Tanganyikabacteria) [5–7]. CPR bacteria are typically ultra-small celled and are predicted to live as obligate episymbionts or parasites of other microorganisms [8–10]. Despite their ubiquity in the microbiomes of diverse aquatic ecosystems, the ecological roles of CPR bacteria and the extent to which their gene content is shaped by environmental factors remain poorly understood. These questions could be addressed in part using genome-resolved metagenomics in systems with strong geochemical gradients.

Here, we examine microbial community composition and function in Lac Pavin, a meromictic lake in central France with a strong oxygen gradient, high methane concentrations in the anoxic water column and sediments, and very low methane concentrations in the upper water layer. We performed genome-resolved metagenomics to obtain hundreds of draft and high quality genomes for members of diverse lineages of bacteria, archaea, and their viruses (phages), and use these genomes to examine changes in C1 metabolic potential across the lake’s depth and oxygen gradient, including methane production and consumption. Together, our analyses shed new light on the diversity and metabolism of Lac Pavin’s microbial communities and how the lake’s geochemical conditions differentially shape the gene inventories of CPR bacteria, other bacteria and archaea, and phages.

## RESULTS

### Diversity of microbial communities across Lac Pavin’s water column and sediments

To examine microbial communities in Lac Pavin, we sampled the water column and shallow sediments of Lac Pavin annually from 2017-2019 (Table S1). The first year of sampling focused on the lake’s oxycline (40-60 meters), whereas subsequent sampling in the second and third years primarily targeted the permanently anoxic monimolimnion (∼60-90 meters) and the shallow sediments at depths between 3 and 58 centimeters (Fig. 1bc, Table S1). We also took one sample from the shallow oxygen maximum (12 meters) as a point of comparison. Water column samples were filtered to the 0.2 - 20 µm size fraction, and whole-community genomic DNA was extracted and sequenced (Materials and Methods). In addition, one post 0.2 µm fraction was used to extend genomic sampling to capture ultra-small cells and phages.

**Figure 1.**
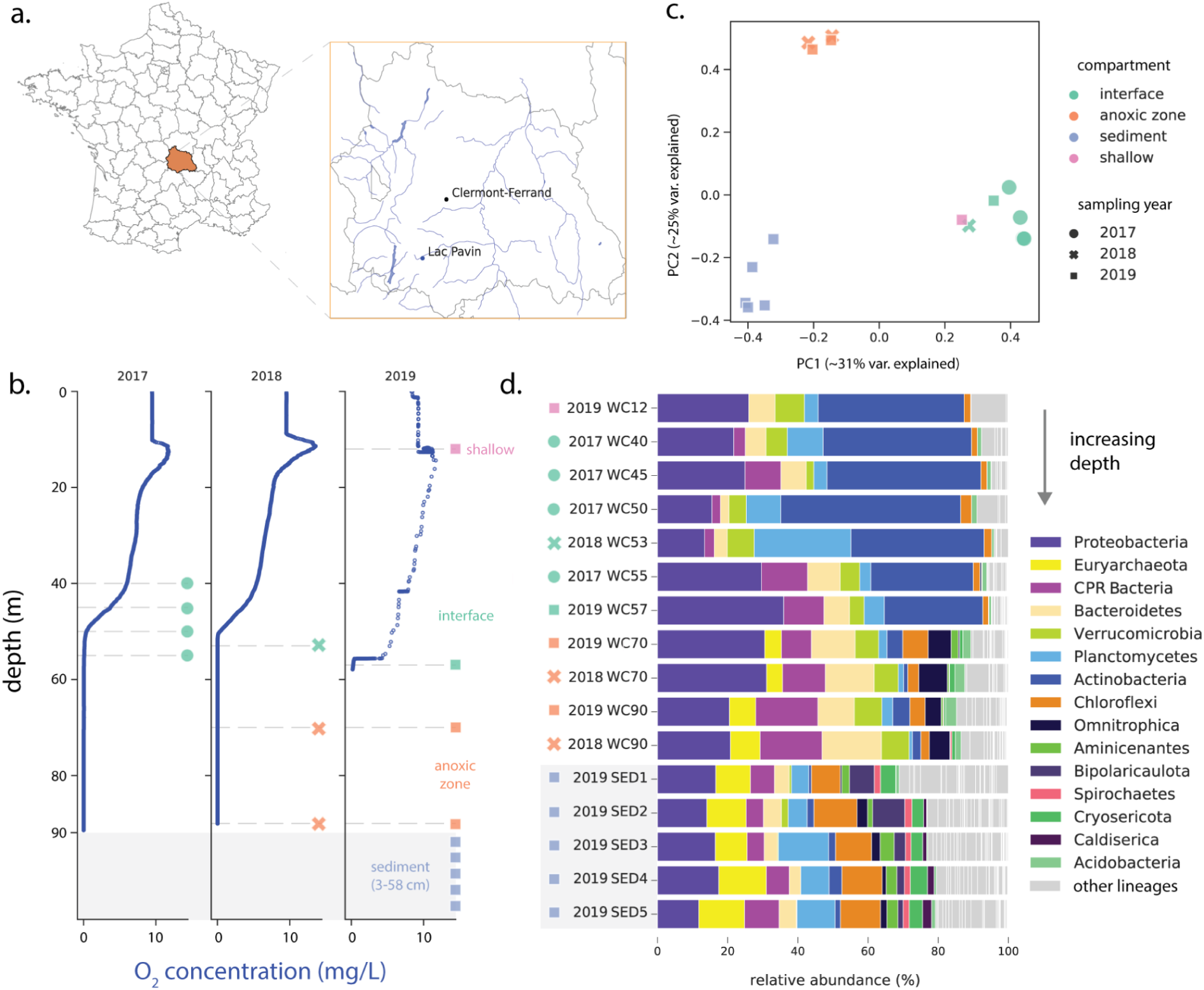
Physical characteristics and microbial communities in Lac Pavin. **a)** Map showing location of the meromictic Lac Pavin in the Auvergne region of central France. **b)** Oxygen profiles across the lake water column during each of the three yearly samplings. Depths sampled on a given year are indicated with a dashed gray line, and colored shapes match those in panels **cd. c)** Principal coordinates analysis of microbial community composition at the ‘species group’ level. **d)** Microbial community composition in Lac Pavin at the phylum level, based on relative coverage of representative scaffolds encoding the ribosomal protein S3. Samples are ordered from shallow to deep, including the five sediment samples collected in 2019. **N.B**. For legibility, only the 15 lineages with the highest median abundance across all samples are shown, all others are indicated in gray as “other”. Abbreviations: PC, principal coordinate; var, variance; CPR, Candidate Phyla Radiation, WC, water column; SED, sediment.

First, we surveyed assembled metagenomic scaffolds for ribosomal protein S3 (rps3) sequences and assigned gene sequences to ‘species groups’ (Materials and Methods). Coverage of these marker genes was then used to examine microbial community similarity across sampling depths/years. A principal coordinates analysis (PCoA) suggested that samples from the same lake compartment, e.g. the samples from the anoxic zone, showed highest similarity to each other across years and were distinct from samples from other compartments, consistent with the permanently stratified nature of the lake (Fig. 1c). Based on community composition, the shallow water column (12 m) most closely resembled samples from the lake interface (Fig. 1c), despite its relatively high oxygen concentration (Fig. 1b).

We investigated the specific phylogenetic groups responsible for driving dissimilarity between microbial communities in the lake, assigning rpS3-defined species groups to phylum-level lineages. Microbial communities from the water column interface were highly enriched in members of the *Actinobacteria* relative to other samples, consistent with findings from other lakes with pronounced oxyclines [7,11] (Fig. 1d, Table S2). The shallow regions of the lake also displayed higher abundances of Cyanobacteria and Planctomycetes compared to the anoxic zone of the water column, where members of the Proteobacteria, Bacteroidetes, and Omnitrophota were particularly abundant (Fig. 1d, Table S2). Finally, the shallow sediments were enriched for members of the Chloroflexi, Cryosericota, Caldiserica, Bipolaricaulota (candidate division OP1), DPANN archaea, and a variety of methanogens, some of which were also found in the anoxic zone (Fig. 1d, Table S2). Surprisingly, the shallowest sediment horizon (3-5 cm) also harbored levels of Cyanobacteria comparable to those in the shallower water column, despite the lack of light (Table S2). One of the three Cyanobacteria species found in the sediment at the times of sampling was also found in the photic zone, suggesting dispersal to the sediment via vertical settling. Notably, members of the Candidate Phyla Radiation were found across the lake’s oxygen gradient and shallow sediments, but were barely detected in the region of the oxygen maximum (Fig. 1d).

### Recovery of draft quality bacterial and archaeal genomes from Lac Pavin

To more closely examine the phylogenetics and metabolic potential of key community members, including CPR bacteria, we generated metagenomic bins by grouping assembled scaffolds into draft genomes with both manual and automated approaches (Materials and Methods). The resulting bins were de-replicated at the species level (95% ANI) and manually curated using Anvi’o and ggKbase tools (Materials and Methods). Refined bins were then filtered again for quality (≥70% completeness) to yield a final set of 738 non-redundant, draft-quality or better genomes for downstream analysis (Table S3). Taxonomy assignment via multiple methods indicated that this set included genomes from most major groups identified by marker gene analysis, including 106 draft genomes from the CPR bacteria, 574 from other bacteria, and 58 from archaea (Table S3).

### Contributions of bacteria and archaea to C1 metabolism over an oxygen gradient

We searched the draft genomes for marker genes for pathways involved in C1 metabolism, carbon fixation, and methane metabolism (Table S4, Materials and Methods) [12]. Using these results, we assigned presence or absence of metabolic functions (pathways) across the set of non-CPR bacteria and archaea and explored their relative abundance in each lake compartment (CPR function and gene content is discussed below). Genomic analyses suggest that the capacity for aerobic carbon monoxide (CO) oxidation is most prevalent at the shallow oxygen maximum and in the interface zone of the lake ecosystem, consistent with the presence of oxygen in both of these lake compartments (Fig 2a). In these regions, CO oxidation was apparently dominated by members of the *Actinobacteria* (Fig. S1), which represented between 3.5 and 18.6% of the microbial community (Table S5). Surprisingly, we also found some evidence for the capacity for aerobic CO oxidation in the anoxic sediments (Fig 2a), primarily in a single genome from the candidate phylum *Bipolaricaulota* (candidate division OP1; Fig. S1). However, the genes encoding CO oxidation were encoded on a very short contig (∼2200 bp), and thus their taxonomic affiliation could not be confidently assigned.

**Figure 2.**
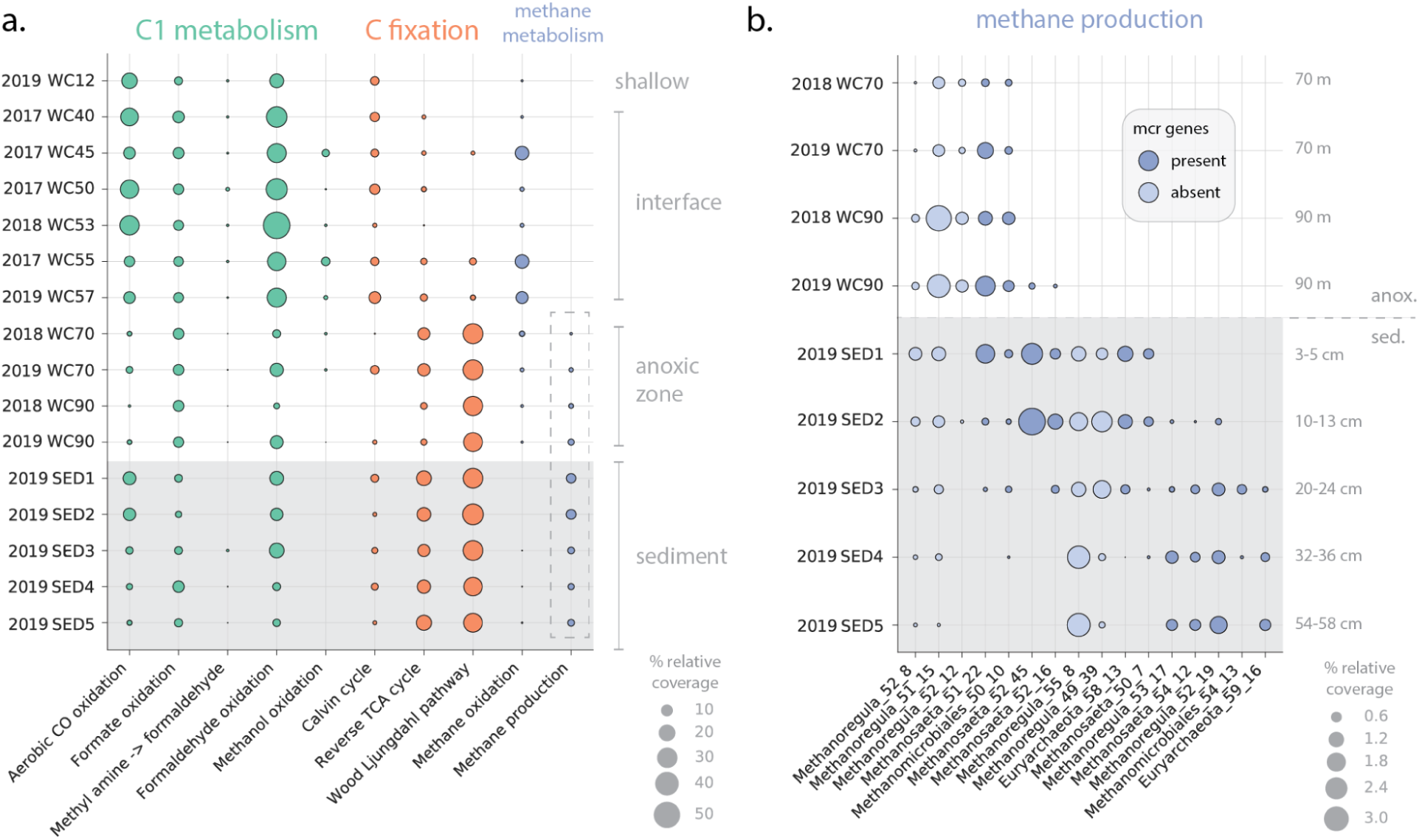
Genetic potential for select carbon metabolism across the water column and sediments of Lac Pavin. **a)** Relative abundance of genomes encoding C1 pathways across metagenomic samples. Bubble size indicates the combined coverage of all organisms encoding a given pathway in each sample normalized by the total coverage of all genomes in the quality-filtered, non-redundant set. Dashed box around methane production indicates the data points expanded in part **b)**, showing the relative abundance (again, as indicated by percent of total genome coverage) for organisms from methanogenic lineages, with and without *mcr* genes. Abbreviations: CO, carbon monoxide; TCA, tricarboxylic acid cycle; anox., anoxic zone; sed., sediment; *mcr*, methyl-coenzyme M reductase.

Biological methane oxidation is thought to act as the major ‘sink’ for methane in Lac Pavin [13]. We thus examined the distribution of genes associated with this process, finding that genomes containing components of the *pmoABC* operon were most abundant in the interface, where in combination they reached a maximum of 13.8% of the microbial community (Fig. 2a, Table S5). Recovered *pmoC* genes were primarily affiliated with the known methane-oxidizing lineage *Methylococcales*, including the genus *Methylobacter*. Some methane oxidizers were present at low abundance in the anoxic zone, suggesting that they may be performing methane oxidation under microaerophilic conditions in this compartment [13,14]. On the contrary, potential methane oxidizers were essentially undetectable in the sediments, where genetic potential for methane production was most concentrated (Fig. 2a). Additionally, we did not detect ANME (anaerobic methanotrophic) archaea capable of anaerobic methane oxidation in either the anoxic zone or the sediments. Thus, overall, we report that methane oxidation is largely spatially separated from methane production in Lac Pavin, consistent with previous amplicon-based studies [14]. A subset of potential methane oxidizers encoded genes for methanol oxidation, which had a parallel distribution through the water column (Fig. 2a, Table S5), suggesting that these bacteria may also be capable of catalyzing the oxidation of methane to formaldehyde.

Microorganisms with the genetic capacity to oxidize formaldehyde were also highly abundant in the lake interface, reaching a maximum relative abundance of ∼51% in one water column sample from 53 meters. Potential for formaldehyde oxidation in the interface was dominated by members of the *Actinobacteria, Planctomycetes*, and *Proteobacteria*, whereas in the sediments *Planctomycetes* and several methanogenic archaea were the greatest contributors. These findings are consistent with the prior discovery of divergent formaldehyde-activating enzymes (*fae*) in *Planctomycetes* [15] as well as the association of fae with methanogenesis [16]. The genetic potential for oxidation of formate, a breakdown product of formaldehyde, was more evenly distributed across the water column (Fig. 2).

Genes encoding the central complex involved in methane production (methyl-coenzyme reductase, alpha and beta subunits, MCR) are primarily localized to the lake sediments, but were also detected in the anoxic zone (Fig. 2). We recovered eleven high-quality, non-redundant archaeal genomes containing the MCR genes that were affiliated with the lineages *Euryarchaeota* (including members of the order *Methanomassiliicoccales)* as well as *Methanosaeta* and *Methanomicrobiales* (including the genus *Methanoregula*), which are commonly detected in freshwater lakes [17]. Read mapping showed that these organisms represent between ∼0.05-3.5% of the microbial community in each sample but are differentially distributed across the deeper water column and sediments (Fig. 2b). Recovered *mcr* genes in the deep water column (70 m) and at the water-sediment interface (90 m) were almost entirely attributable to two organisms (one *Methanosaeta* and one *Methanomicrobiales* organism). The potential methane producers in the sediment were more diverse at the genus level, particularly in the shallower sediments (3-24 cm) (Fig. 2b). *Mcr* genes in the deeper sediments (32-58 cm) were primarily attributable to *Methanoregula, Methanosaeta*, and *Euryarchaeota* genomes which were rare or non-existent in the shallower sediments as well as the water column (Fig. 2d). Five widespread genomes affiliated with the *Methanoregula* were also recovered by our binning efforts; however, they did not contain the *mcr* gene cluster, potentially due to genome incompleteness (Fig. 2b).

We also examined the genetic potential for carbon fixation across the water column and sediments. The Calvin Cycle (form I and II RuBisCO) was present throughout the water column, but was most prevalent in the shallower lake zones (Fig. 2a), primarily among Cyanobacteria, Actinobacteria, and Proteobacteria (Fig. S1). In contrast, we found that the genetic potential for CO_2_ fixation via the Wood-Ljungdahl (W-L) pathway was nearly absent in oxygenated parts of the lake (Fig. 2a) but was common among Chloroflexi, Proteobacteria, and Euryarchaota in the anoxic water column and sediments, which together comprised ∼25-31% of the community (Fig. S1, Table S5). To a lesser extent, organisms with the genetic potential for the reverse TCA cycle were also associated with the lake’s anoxic zone and sediments, with the exception of samples taken from 90 meters at the anoxic/sediment interface (Fig. 2a), where this capacity was rare. We found no evidence for alternative carbon fixation pathways involving 3-hydroxypropionate cycle or 4-hydroxybutyrate among Lac Pavin organisms.

Together, these findings point to depth/oxygen as a structuring force for not only microbial community composition in Lac Pavin, but also for certain metabolic capacities related to carbon cycling that display clear evidence of spatial stratification.

### CPR bacteria with the metabolic potential to incorporate CO_2_ via RuBisCO

Previous work based on 16S rRNA amplicon-based approaches showed that Lac Pavin hosts diverse communities of bacteria from groups now recognized to be part of the CPR, and showed that these communities vary with depth [5]. Given prior research demonstrating the diversity of RuBisCO enzymes among these organisms [18,19], we hypothesized that some CPR in Lac Pavin may play a role in carbon cycling via CO_2_ incorporation into biomass. We identified form II/III RuBisCO in members of the Absconditabacteria and the Komeilibacteria (Parcubacteria radiation) and form III-c in another Komeilibacteria (Fig. 3bcd), a lineage in which form III-c has not previously been observed. Both form II/III and all form III RuBisCO likely function by incorporating carbon dioxide with a free nucleotide to form glyceraldehyde-3-phosphate, a central glycolytic intermediate [20].

**Figure 3.**
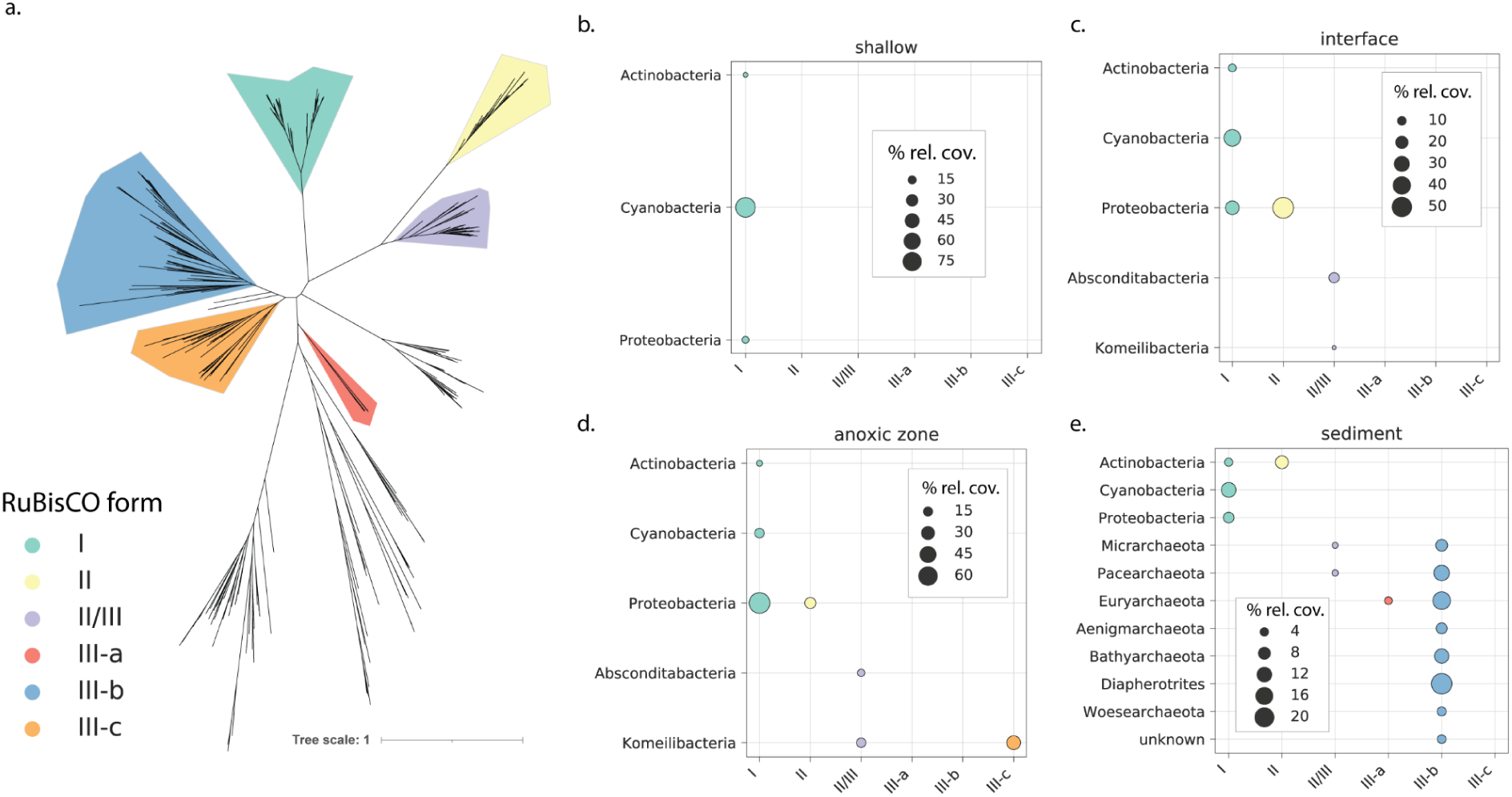
Diversity and distribution of RuBisCO in Lac Pavin. **a)** Unrooted phylogenetic tree showing relationships among major clades within the RuBisCO superfamily. **b-e)** Distribution of community metabolic potential for carbon fixation/CO_2_ incorporation via RuBisCO in each lake compartment. Bubble size represents the percentage of coverage for all scaffolds with RuBisCO attributable to a given lineage/RuBisCO pair. This percent coverage is averaged across all samples within a lake compartment.

We quantified the relative abundance of CPR RuBisCOs in comparison to other RuBisCO genes encoded by other bacteria and archaea, including those found both in genomic bins as well as in unbinned scaffolds. In the interface, form II/III from the Absconditabacteria composed on average 13.6% (ranging from ∼10-17%) of RuBisCO gene copies and form II/III from the Komeilibacteria contributed only ∼2.2% of gene copies on average (Table S6). However, in the anoxic zone, Komeilibacteria accounted for an average of 32.6% of RuBisCO gene copies, yet this varied dramatically, ranging from 1% to 83% across samples (Table S6). Interestingly, within the anoxic zone just above the sediment-water interface (90 meter sample), RuBisCO from Komeilibacteria was the largest contributor to RuBisCO genetic potential (Fig. 3c). CPR bacteria with RuBisCO were barely detectable in the lake sediments, where community potential for CO_2_ incorporation via RuBisCO was dominated by archaea bearing the form III-b enzyme (Fig 3d, Table S6). The most abundant genes were encoded by various DPANN lineages, including the Pacearchaeota and Diapherotrites (Fig. 3d).

### Diversity and distribution of CPR bacteria in Lac Pavin

We examined the broader phylogenetic diversity of CPR bacteria in Lac Pavin using genome sequences from which we extracted and concatenated 16 syntenic, ribosomal proteins. This analysis revealed that CPR bacteria from Lac Pavin are highly diverse and include members from nearly all groups currently recognized within the CPR. Beyond the Absconditabacteria and Komeilibacteria mentioned above, we also identified organisms from the Saccharibacteria, Peregrinibacteria, Microgenomates, and recently delineated clades of the Parcubacteria [21], each of which each contains numerous sub-lineages (Fig. 4a).

**Figure 4.**
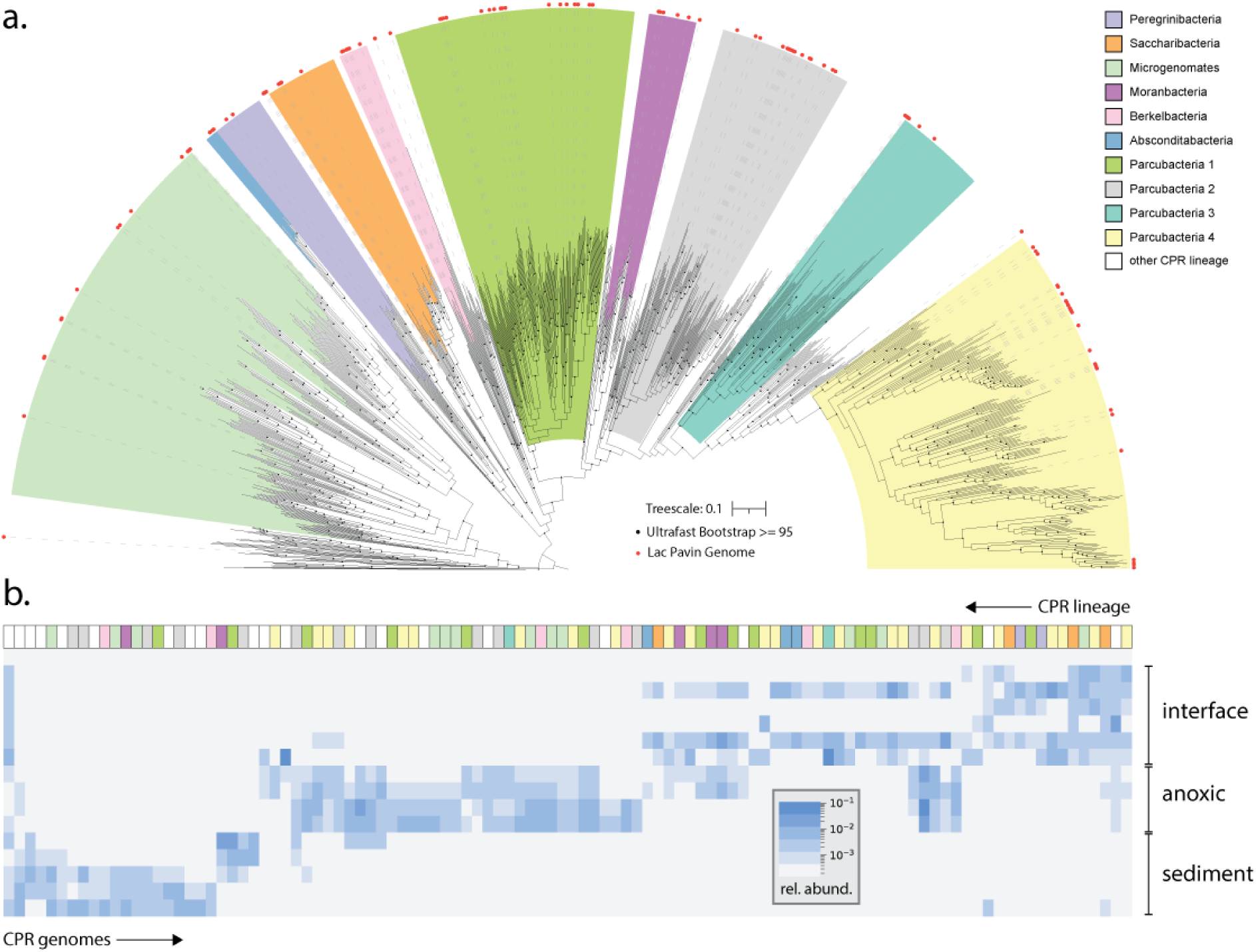
Phylogenomics and abundance distribution of CPR bacteria in Lac Pavin. **a)** Phylogenetic placement of CPR bacteria from Lac Pavin (red circles) using a set of reference genomes from across habitats. Major phylogenetic groupings are indicated; however, note that not all are equivalent in terms of rank. Scale bar represents the average number of substitutions per site. **b)** Relative abundance (coverage) of individual CPR genomes across samples. Lineage assignment (matching the color scheme from panel **a**) is indicated at the top of panel **b**.

By mapping metagenomic reads back to the draft CPR genomes, we determined the abundance of individual CPR organisms across the lake’s oxygen gradient. While CPR bacteria were detectable throughout the water column and shallow sediments, they were most abundant in the lower interface and anoxic zone (55-90 meters depth) (Fig. 1d). Intriguingly, different species from the same lineages populated each zone, e.g., different species of Berkelbacteria were uniquely associated with the interface, anoxic zone and sediments (Fig. 4b, Table S7). In contrast, some lineages were only detected in a single compartment: for example, members of the Saccharibacteria were mostly restricted to the lake interface (Fig. 4b, Table S7).

### Broad patterns of gene content in CPR bacteria

To complement our analyses of RuBisCO in CPR, we examined the extent to which entire predicted proteomes of CPR bacteria vary throughout Lac Pavin’s water column. Using a protein clustering approach that is agnostic to gene function [22], we resolved ∼1500 protein families (≥ 5 proteins in size) among all CPR bacteria and visualized their distribution within the draft genome set. Clustering based on protein family presence/absence revealed a large ‘module’ (or group of protein families) that were found in nearly all CPR genomes (Fig. S2), paralleling results from a prior analysis of CPR proteomes [22]. As expected, this ‘core’ module contained key functions such as transcription, translation, and cell division. Outside of this core module, most other protein families were patchily distributed (Fig. S2).

While there was some evidence for groups of protein families that are specific to a small number of CPR genomes (narrow, horizontal strips in Fig. S2), no large modules were associated with organisms from the interface, anoxic zone, sediment, or combinations thereof. Detailed statistical analysis revealed only 22 protein families moderately to highly enriched in a specific compartment (enrichment ratio ≥3, p≤0.05, two-sided Fisher’s exact test corrected for multiple comparisons). Most of these families (∼82%) were enriched in CPR bacteria from the interface, and included protein families that were involved in the metabolism of malate (K00027), zinc transport (K09815), and metabolism of certain amino acids (K00928) (Table S8). Also notable was a family confidently annotated as an Fe-Mn superoxide dismutase (K04564) that was moderately enriched (enrichment ratio ∼2.7) in CPR bacteria from the interface relative to those from the anoxic zone and sediments (Table S8). Overall, these results suggest that while organisms from the interface may have slight differences in their metabolic capacities and stress responses compared to those from the anoxic zone and sediments, in general proteomes of CPR bacteria across the water column are not highly differentiated from one another.

### Diversity and characteristics of recovered phage genomes

Viruses of bacteria (bacteriophages, or phages) are important components of the ‘microbial loop’ in the shallow regions of Lac Pavin and its associated sediments, where they are known to exhibit significant morphological and genomic diversity [23–25]. Here, we leveraged our metagenomic data to extend analyses of the diversity, distribution, and metabolism of Lac Pavin phages that were retained by our sampling/filtration methods (Materials and Methods). We refined a preliminary set of 17,151 potential phage scaffolds to yield a final set of 767 non-redundant scaffolds (focusing on those larger phages with ≥50 kb genome size) that could be confidently identified as phages or bacterial/archaeal genome fragments with integrated prophages (Materials and Methods, Table S9). Approximately 7.7% of phage scaffolds were circularized and thus likely represent complete genome sequences (Table S9). We merged these newly-reported phage scaffolds with those from our previous studies of the same environment [26,27] and subjected the total set of ∼800 non-redundant phage scaffolds to a network analysis based on protein content.

This analysis revealed that many phages from Lac Pavin fall into broad clusters with known phages from diverse environments, suggesting they are part of well-sampled clades defined by protein content (“mixed” clusters, Fig. 5a). Of particular interest were Lac Pavin phage scaffolds that clustered with megaphages from diverse environments [26], phages encoding the S21 ribosomal protein from freshwater environments [27], and phages from another large freshwater lake in Japan, Lake Biwa [28] (Fig. 5b). We also recovered 52 phage clusters that contained only newly-reported sequences from Lac Pavin (Fig. 5a). These clusters were generally small (5 or fewer related sequences) and contained similar or higher sequence diversity (as measured by pairwise distances) than clusters of the same size that included both Lac Pavin and reference sequences (“mixed”) or only reference sequences (Fig. 5a). However, the majority of Lac Pavin phage scaffolds (n=477) did not cluster with any other phages (cluster size = 1, Fig. 5a). These results suggest that Lac Pavin viromes may also harbor considerable genomic novelty in addition to members of broadly distributed, well-sampled viral groups.

**Figure 5.**
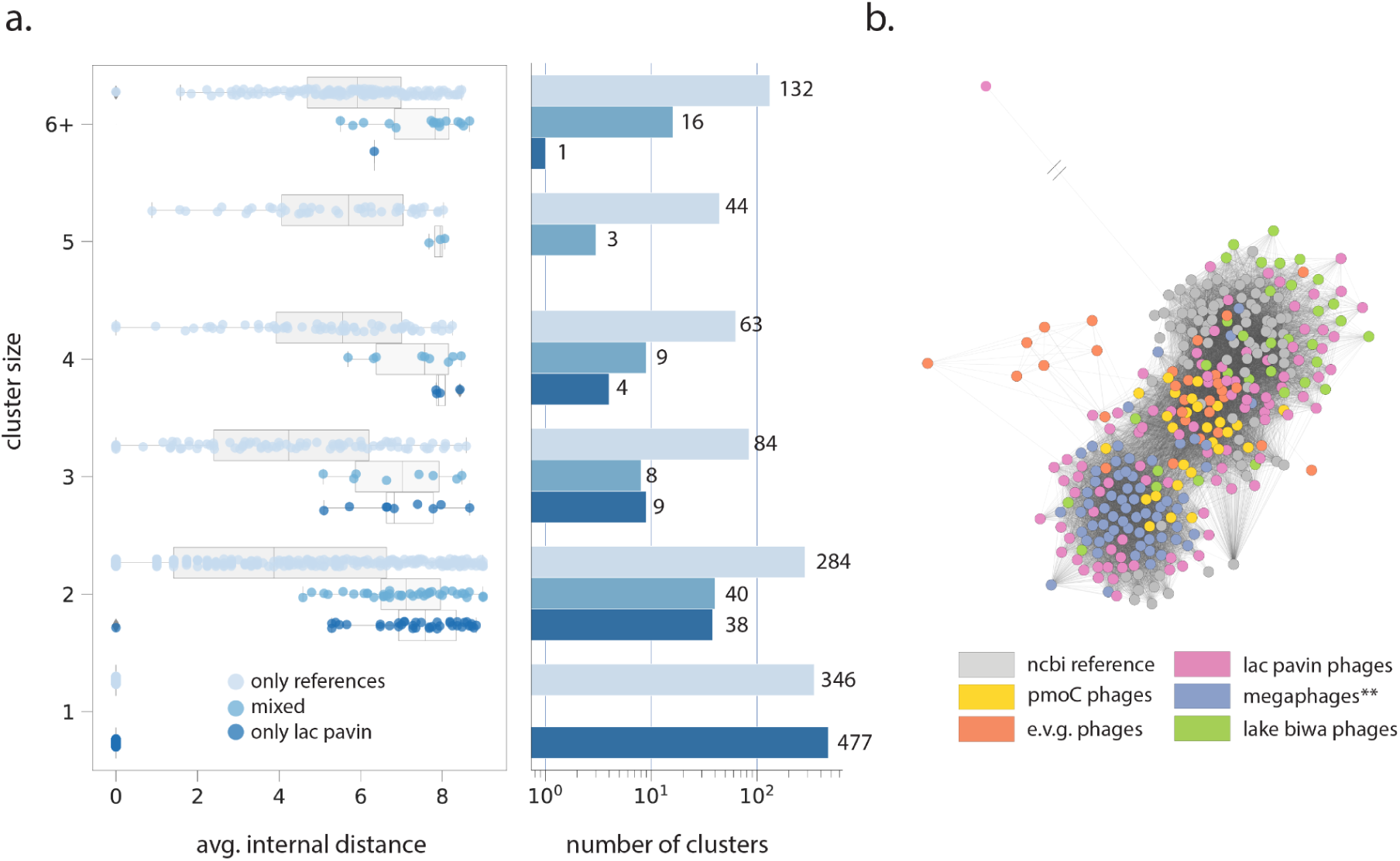
Diversity of phages (≥50 kb genome size) in Lac Pavin. **a)** Cluster composition, sequence diversity (described by average pairwise distance among nodes within the cluster), and size for a network analysis describing the relationship of Lac Pavin phages to known references based on protein content. “Mixed” clusters are those containing both reference and Lac Pavin phage genomes. **b)** Subset of phage network analysis displaying interconnectedness between Lac Pavin phages and those from various reference databases. Shorter edges between nodes (phages) indicate a higher degree of similarity based on protein content. **Megaphages from diverse environments (Al-Shayeb, Sachdeva, et al., 2020).

### Phage distribution and metabolism

Motivated by the recent recognition of the importance of large phages across ecosystems, including Lac Pavin [26], we further analyzed the distribution of phages with genomes ≥100 kbp in length across the lake compartments using read mapping. Our results indicate that large phages are present throughout the water column and shallow sediments (Fig 6a, Table S10). While many phages were restricted to certain compartments - e.g., the interface or sediments, we also found evidence for phages that were present in multiple zones (Fig. 6a, Table S10). The three largest megaphages, attaining ∼735 kb (the largest phage genome yet reconstructed) [26], 642 kb, and 623 kb in length were present in the interface and shallow sediment, the interface, and sediment, respectively (Fig. 6a). The largest two of these phages were predicted to infect members of the Bacteroidetes (Table S9). Other phages predicted to infect Proteobacteria, Planctomycetes, and Spirochaetes bacteria were detected through the water column and sediments. Surprisingly, one phage from the sediment was predicted to infect a member of the Candidatus Thorarchaeota based on taxonomic profiling of its predicted proteins (Material and Methods). However, given that we did not recover strong evidence for the host archaeon itself in the sediments, the strength of this phage-host association is unclear.

**Figure 6.**
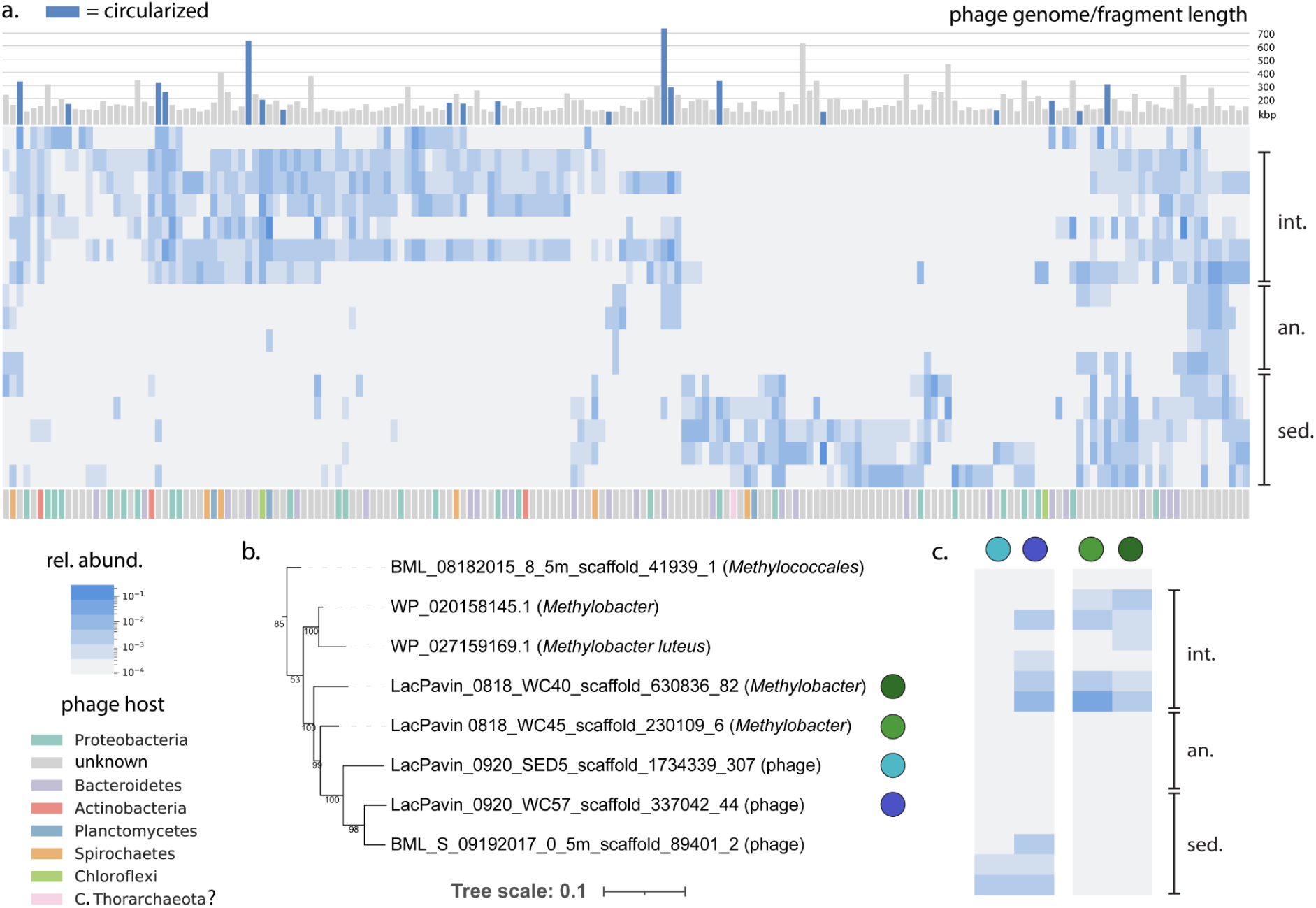
Distribution and metabolism of phages in Lac Pavin. **a)** Relative abundance of large phages (≥100 kbp) based on read mapping. Barplots above the heatmap represent fragment/genome size in kbp). Bars in blue represent phages that are likely to be circularized. Color strip below the heatmap indicates host assignment based on CRISPR-Cas targeting or taxonomic profiling of phage genes. **b)** *pmoC* protein phylogeny representing the evolutionary relationships between phage sequences and sequences from potential *Methylobacter* hosts. **c)** Relative abundance mapping of phages encoding *pmoC* (left) and potential hosts (right) through the water column and shallow sediments. Abbreviations: int., interface; an., anoxic zone; sed., sediments; rel., relative.

The extent to which environmental factors determine the gene content of phages is unclear. We tested whether any protein clusters derived from our phage genomes displayed signatures of enrichment in particular compartments of the lake. Paralleling our findings for CPR bacteria, we did not observe strong evidence for gene content differentiation by depth, with only two protein families statistically enriched in phages from the interface and anoxic zones, respectively (p≤0.05, two-sided Fisher’s exact test corrected for multiple comparisons) (Table S11). While not statistically enriched in any compartment, we also found two phages encoding a *pmoC* gene. These genes have been previously reported in diverse phages from freshwater lakes [29] potentially implicating them in Lac Pavin’s cycle of methane production (primarily in the sediments) and methane oxidation (primarily in the water column interface). While we were not able to obtain host assignments for these phages by CRISPR-Cas targeting or by taxonomic profiling, we observed that the phage *pmoC* sequences were closely related to those from *Methylobacter* genomes resolved from the Lac Pavin water column interface (Fig. 6b, Table S12), suggesting them as possible hosts. Surprisingly, relative abundance mapping showed that both phages harboring a *pmoC* gene were found in the deeper sediments, where potential *Methylobacter* hosts were not detected (Fig. 6c), suggesting either an alternate host in these samples or burial and preservation of phage particles in the sediments. Supporting the second explanation, we note many interface-associated phages are absent in the anoxic zone but present in the sediment (Fig. 6a). One of the *pmoC* phages was also found in the water column interface alongside *Methylobacter*, consistent with a possible host relationship (Fig. 6c).

## DISCUSSION

Despite differences in morphology and origin, meromictic lakes generally have similar physical characteristics that shape their microbial communities [30]. Our results suggest that like other similar lakes, the availability of oxygen across the water column and shallow sediments of Lac Pavin shapes both microbial community composition and their metabolic capacities. We provide high-quality genomic data for numerous key lineages in this ecosystem, including abundant members of the *Actinobacteria*, which dominate in the shallow water column of Pavin and other similar lakes [31,32]. On the other hand, we found very little if any evidence for the photosynthetic purple/green sulfur bacteria that frequently occur in the oxycline of meromictic lakes, sometimes at extremely high densities [33–35]. While the absence of these bacteria may be due to relatively low light intensities in Lac Pavin’s oxycline compared to those in other lakes, there is sufficient light to support some members of the *Cyanobacteria* in this zone (Table S2). Another factor may be the low abundance of sulfide in this zone of the water column.

Microbial metabolism is a major driver of biogeochemical cycling within lake ecosystems. We examined the extent to which changes in community composition across depth in Lac Pavin are accompanied by changes in metabolic potential, with a specific focus on a subset of C1 carbon metabolic pathways. Our analyses show that oxygen impacts metabolic potential, in part by selecting for bacteria and archaea with certain C1 and carbon fixation pathways in different zones of the lake. In particular, the Wood-Ljungdahl and reverse TCA cycle showed strong signals of stratification and were essentially restricted to the lake’s anoxic zone, consistent with findings from another stratified lake [36], although that lake is ∼15 times shallower. Thus, potential for autotrophy via these pathways may vary predictably with geochemical conditions in a scale-independent manner. We found only minimal evidence for anaerobic methane oxidation (e.g., by ANME archaea) and thus expand upon prior evidence for spatial segregation of methane production and consumption in this lake. At a finer scale, we leveraged our genome-based approach to determine that the species composition of methane producers changes on the scale of centimeters within the shallow sediments (Fig. 2b). This observation hints at the possibility of niche partitioning among related methanogenic lineages, a trend which may continue into the deeper sediments not sampled in this study [37].

A prior study used amplicon sequencing to document CPR in Lac Pavin [5]. Here, we build upon those results by providing high-quality draft genomes for over 100 CPR bacteria, showing that Lac Pavin harbors a high abundance of CPR bacteria with representatives from virtually all of the major lineages. Broadly speaking, gene content in CPR bacteria, even within specific lineages, can be highly variable [21,22] raising the important question as to what governs gene content in these organisms. One possibility is that ecosystem characteristics, like local geochemistry, drive the distribution of capacities in CPR bacteria. On the other hand, metabolic capacities of the host cell(s), and/or the spectrum of resources that are exchanged between them, may determine which pathways are present in a given CPR organism.

Here, we found that CPR bacteria display relatively little evidence for metabolic differentiation across depth in Lac Pavin, despite large variations in geochemical conditions. This suggests that within certain habitat types, CPR gene content is likely primarily determined by the metabolic capacities of their hosts, rather than geochemistry. In Lac Pavin, this appears to stand in contrast with patterns of gene content observed for other, non-CPR organisms, which are likely shaped more heavily by the availability of oxygen, light, and other key nutrients. Synthesizing these observations, we postulate that geochemistry selects indirectly for CPR bacterial types via selecting for hosts with metabolisms that utilize the spectrum of resources provided across the water column.

One exception to the weak linkage between geochemistry and CPR metabolic traits is the moderate enrichment of superoxide dismutases in CPR bacteria from the partially aerobic interface relative to those from the anoxic water column and sediments. Proteins from this family may be involved in radical detoxification at the lake interface where oxygen levels are relatively high compared to deeper compartments (Fig. 1b). Other analyses have shown that superoxide dismutases are rare in CPR from anoxic human oral microbiomes [38], yet they do occur in organisms from oxic groundwaters [39]. Taken together, these findings suggest that this protein family may be one of the few components of CPR genomic repertoires that may correlate with environmental conditions across ecosystems.

Despite their reduced metabolic platforms, CPR bacteria probably still impact biogeochemistry throughout lakes. Along with genes that shape carbon, sulfur, and nitrogen cycles [40–42], RuBisCO is encoded by some CPR bacteria and could be used to fix carbon dioxide at some rate [18–20]. Our analyses of RuBisCO in Lac Pavin show that some CPR bacteria carry form III variants in the water column, but RuBisCO genes were not found in sediment-associated CPR. There, DPANN and other archaea tend to carry form III RuBisCO, suggesting spatial segregation of the organisms whose genomes encode this enzyme form. Given their wide environmental prevalence and the relatively little biochemical research on form III RuBisCOs [43], additional work is required to determine their rates of carbon dioxide incorporation. With this information it will be possible to quantify the potential contributions of form III RuBisCOs to carbon fixation *in situ*, including by CPR bacteria.

Phages, including the largest megaphages yet reported, were distributed throughout the water column and sediments in Lac Pavin, but like CPR bacteria, their metabolic capacities did not vary significantly by depth (Fig. 6, Table S10-11). This observation is consistent with the notion that gene content in phages is primarily selected for by interactions with their microbial hosts. This selection may extend to recently described metabolic genes in phages that may augment host metabolism of the same or similar compounds [44–47], like the phage *pmoC* genes described above (Fig. 6bc). Taken together, our findings raise the intriguing possibility that selection on CPR gene content may resemble that on phages more so than that shaping those of other bacteria, in part due to their shared strong reliance on hosts. However, there is currently no strong evidence for recent lateral transfer between CPR bacteria and known hosts validated by laboratory co-cultivation [48], as is known for phages. This suggests the evolutionary processes that shape gene selection in CPR bacteria and phages probably differ.

An final important feature of our study is the concurrent investigation of microbial communities in sediments in tandem with those in the overlying lake water column. In addition to finding functional partitioning likely driven by geochemical conditions, we also detected members of the Cyanobacteria, *pmoC* phages without their expected methanotrophic hosts, and phages from the interface zone that were present in the sediment. It is perhaps unsurprising that sediments can accumulate cells via settling; however, our findings raise the possibility that phage particles are preserved there and thus are capable of contributing to long term carbon storage. Overall, our analyses provide insights into the factors shaping the gene content of CPR bacteria, non-CPR bacteria, archaea, and phages through a stratified lake, revealing the differential impact of geochemistry on their genetic repertoires.

## AVAILABILITY OF DATA AND MATERIALS

Read data and draft genomes from this study are available through NCBI at PRJNA852163. Genome accession information for bacterial/archaeal genomes is also listed in Table S3. Custom codes for the described analyses are available on GitHub (https://github.com/alexanderjaffe/lac-pavin-metagenomics). All supplementary figures, tables, and genomes (including phage/viral genomes) are also available through Zenodo (https://doi.org/10.5281/zenodo.6857590).

## ACKNOWLEDGMENTS

We thank Yue Clare Lou, Spencer Diamond, Shufei Lei, Adair Borges, Marie Schoelmerich, Lin-Xing Chen, Alex Thomas, and Rohan Sachdeva for helpful discussions and informatic support. We also thank Guillaume Borrel and François Enault for their comments on the manuscript, and Lily Law, Christopher Lefevre, and Didier Jézéquel for their assistance with sampling logistics. Funding was provided by the Berkeley Fellowship to A.L.J. and Moore Foundation Grant 71785 to J.F.B. We also thank the Innovative Genomics Institute at UC Berkeley, Y. Justin Choi, Shana McDevitt, and QB3 Genomics at UC Berkeley. Sequencing was partially supported by NIH S10 OD018174 Instrumentation Grant.

## AUTHOR CONTRIBUTIONS

A.L.J, C.J.C., A.C.L., and J.F.B. designed the project. A.L.J., J.L., and J.F.B. performed bioinformatic analyses and genome curation. A.L.J, C.J.C., C.B., H.B., J.C., F.P., A.H.L.J., and A.C.L. performed sample collection. C.B., H.B., J.C., F.P., A.H.L.J., and A.C.L. performed sample preparation and DNA extractions. A.L.J. and J.F.B. wrote the manuscript. All authors made comments on the manuscript.

## MATERIALS AND METHODS

### Sample collection and measurements of oxygen

Prior to sampling, oxygen concentration and saturation were measured through the water column with a SDOT dissolved oxygen data logger (NKE Instrumentation). All water column samples were collected using a 12 liter Niskin bottle. Between 3 and 5 bottles were taken for each water column sample (36-60 liters total) and all the water from one depth was pooled and filtered by tangential flow filtration (0.2 µm cartridge) to yield concentrated per sample. The 12 meter and 57 meter samples from 2019 were pre-filtered before tangential flow filtration with a 20 µm filtration tissue. Filtrate from the 2018 70 meter sample was subsequently re-filtered using a 30 kilodalton cartridge to yield a small size fraction concentrate (sample 70S, < 0.2µdDa). Filtrate from each depth was then split into fractions, each of which was subjected to centrifugation at 24,500g (30 minutes, 4°C). Supernatant was removed and pellets from each fraction from the same samples were resuspended in TE buffer (Tris 10mM; EDTA 1mM; pH8) and pooled. The pooled samples were centrifuged one final time as described above and pellets were stored at −20°C until DNA extraction.

In 2019, a sediment core was taken and 5 strata were collected: 3 to 5 cm (SED1), 10 to 13 cm (SED2), 20 to 24 cm (SED3), 32 to 36 cm (SED4), and 54 to 58 cm (SED5). Each stratum was homogenized and stored at −20°°C for DNA extraction. All sample metadata is listed in Table S1.

### DNA extraction and sequencing

DNA was extracted according to the method described by [49], with slight modifications. Each pellet was defrosted and incubated in TE buffer (pH 8). Cells were lysed with lysozyme (Sigma, final concentration 20 mg/ml) for 30 min at 37°C and then incubated for 1 hr at 37°C with sodium dodecyl sulphate (SDS, final concentration 0.5%), proteinase K (Sigma, final concentration 0.1 mg/ml) and RNaseA (Sigma, final concentration 0.8 μg/ml). Cell debris, proteins and humic substances were removed by adding 100 μl of a solution of cetyltrimethylammonium bromide (CTAB)/NaCl (10%, w/v, and 0.7 M, respectively) and 80 μl of NaCl 5 M for 10 min at 65°C. CTAB-protein/polysaccharide complexes were extracted with 1 volume of phenol/chloroform/isoamyl alcohol (24:24:1) after centrifugation (10,000g, 4°C, 30 min).The aqueous phase was re-extracted with 1 volume of chloroform/isoamyl alcohol (24:1) after centrifugation (10,000g, 4°C, 30 min). Nucleic acids (aqueous phase) were precipitated with 0.6 volume of ice-cold isopropanol overnight and centrifuged (10,000g, 4°C, 30 min). Pellets were washed with 70% ethanol and centrifuged (10,000g, 4°C, 30 min). pellets (nucleic acid) were air-dried and dissolved overnight in 50 μl of H2O. DNA was quantified using the Qubit® 2 .0 fluorometer (Life Technologies) and stored at −20°C until processing.

250mg of sediment from each stratum was defrosted and centrifuged for 30 seconds at 10,000g to remove the supernatant. For the lysis step, samples were heated to 70°C and then vortexed for 10 minutes. DNA was then extracted using the Qiagen DNEasy Power soil kit following manufacturer instructions.

Library preparation and metagenomic sequencing were performed at the QB3 (University of California, Berkeley) Functional Genomics Laboratory / Vincent J. Coates Genomics Sequencing Laboratory. Libraries were sequenced with 150 base-pair, paired-end reads on either a Illumina HiSeq 4000 platform (2017 and 2018 samples) or a NovaSeq S4 platform (2019 samples) with a target depth of 15-25 gigabasepairs per sample.

### Metagenomic assembly, coverage calculation, gene prediction, and preliminary annotation

Metagenomic assembly was performed following the procedures of He et al. 2021 [50]. Briefly, sequencing reads were rid of Illumina adapters using BBTools and were trimmed using Sickle [51] (default thresholds). Quality-filtered reads were then assembled using MEGAHIT (v. 1.2.9, default parameters) [52] and scaffolded using IDBA-UD [53]. Reads were mapped back to the entire assembly for each sample using bowtie2 [54] to compute scaffold coverage values. Assembled scaffolds were subsetted to those ≥1000 bp for gene prediction and binning. Genes were predicted using Prodigal (*meta* mode) [55] and predicted proteins were annotated using USEARCH against the KEGG, UniRef, and UniProt databases.

### Genome binning, refinement, taxonomic assignment, and relative abundance calculation

Genome binning was performed using both a manual and an automated approach. For the manual approach, scaffold information was loaded into ggKbase (ggkbase.berkeley.edu) and scaffolds were binned on the basis of coverage, GC content, taxonomic affiliation, and inventories of bacterial ‘single copy’ genes. For the automated approach, reads were cross-mapped against every other assembly within each biome type using bowtie2. Sediment and water column samples were binned separately. Coverage tables were generated using the jgi_summarize_bam_contig_depths script (bitbucket.org/berkeleylab/metabat/src/master) and passed to MetaBAT2 (minimum contig size 1500 bp) [56] for automated bin generation. Bins derived from the manual and automated approaches were reconciled with DAS Tool [57] to create the best merged set of bins for downstream analysis.

To prepare for the bin refinement step, preliminary taxonomic classifications were performed using GTDB-Tk [58]. Bins associated with the Candidate Phyla Radiation bacteria or DPANN archaea were separated and profiled for reduced sets of marker genes sensitive to lineage-specific losses in these groups [59]. Completeness and redundancy were calculated as the percentage of marker genes present and duplicated, respectively. These quality metrics were combined with those for all other, non-CPR and non-DPANN bins estimated by CheckM [60]. Bins were then filtered to those with ≥70 completeness and de-replicated at 95% average nucleotide identity (ANI) using dRep to create a secondary set for manual curation. All quality-filtered bins were loaded into Anvi’o [61] and visualized individually using the *anvi-refine* command. Bins were refined by removing sets of scaffolds with aberrant coverage profiles across all cross-mapped samples. Completeness and redundancy metrics were also considered. Refined bins were then re-assessed for quality (as above) and additionally profiled with GUNC [62] as an alternative metric of contamination.

Draft genomes from the CPR bacteria and DPANN archaea were classified by extracting 16 (or 14, in the case of DPANN) phylogenetically informative, syntenic ribosomal proteins [63] from each genome (where present) and combining them with comprehensive set of reference sequences from previous studies [21,64]. Each marker protein was aligned individually using MAFFT [65] and trimmed using BMGE (*-m BLOSUM30*) [66]. All individual protein alignments were concatenated to generate a supermatrix. Any organisms for which less than 7 of the 16 marker proteins were present were discarded. Finally, a maximum-likelihood phylogenetic tree was inferred using IQTREE (*-m TEST --b 1000*) [67]. Taxonomic assignments of newly-recovered draft genomes from Lac Pavin were manually assigned using the tree topology and taxonomy of proximal reference sequences. For all other bacteria and archaea, phylum-level taxonomy was assigned using a combination of GTDB-Tk results, protein annotations from ggKbase, and manual BLAST searches of the S3 or L6 ribosomal proteins.

Finally, reads from all samples were re-mapped to the refined bin set using bowtie2. Bin coverage was analyzed using CoverM (https://github.com/wwood/CoverM) with a 95% identity threshold for read mapping and 50% breadth threshold (fraction of genome covered by reads). Relative coverage values for each genome were computed by dividing its mean coverage by the total, summed coverage of all genomes present within a given sample.

### Marker gene-based analyses of community composition

Predicted proteins from all assemblies were probed using a custom Hidden Markov Model (HMM) describing ribosomal protein S3 sequences (rps3) from diverse bacteria and archaea and thresholded using an HMM alignment score of 40 [68]. Recovered rps3 sequences from across samples were subsequently clustered into ‘species groups’ using usearch -cluster_fast (*-sort length -id 0*.*99 -maxrejects 0 -maxaccepts 0*) [69]. For each species group, the longest rps3 protein sequence and the longest scaffold from any species group member were chosen as representatives [68]. Representative protein sequences fragmented into multiple pieces by assembly errors were removed by selecting the longer partial sequence, and all representative sequences smaller than 1 standard deviation below the mean sequence length were removed.

Taxonomic affiliation was assigned at the phylum level using three separate pieces of evidence: the taxonomy of the bin containing the representative scaffold (where relevant), the taxonomy of the top UniRef100 BLAST hit to the representative protein sequence, and the consensus taxonomy of the representative scaffold. Consensus scaffold taxonomy was computed by comparing each predicted protein against UniRef100 using a custom DIAMOND database (diamond blastp) [70]. Hits were filtered to those with 70% or greater coverage of the query sequence and an e-value less than or equal to 1×10^−10^, as done previously [71], and the most common phylum-level designation was taken. If the same phylum-level designation was predicted by any two pieces of evidence, it was assigned as the final taxonomy; otherwise, scaffold taxonomy was manually curated using BLAST searches and annotation information.

Finally, relative abundance of species groups were assessed by mapping all raw sequencing reads to the combined scaffold set using Bowtie2 (default parameters). Coverage was calculated using CoverM with a stringent mapping threshold (99% read identity) and a minimum coverage breadth of 50% of representative scaffolds. Relative coverage of each species group was calculated by dividing its mean coverage value by the total coverage of all species groups in a given sample. To examine overall similarity in community composition across samples, the relative coverage metrics computed above were subsequently used to generate a Bray-Curtis distance matrix (skbio’s diversity package in Python) and subjected to a principal coordinates analysis (implemented in skbio’s stats module).

### Analysis of community-wide carbon metabolism

Proteins were re-predicted for all refined bins using Prodigal (*single* mode). All proteins that were not from Candidate Phyla Radiation bacteria were subjected to annotation with kofamscan [72]. Protein annotations were filtered first by e-value (≤1e-6) and the highest scoring hit per protein was taken. Annotations were searched for a curated, previously published set of hidden Markov models (HMMs) representing key marker genes for pathways involved in C1 metabolism, carbon fixation, and methane metabolism (Table S4) [12]. We next manually adjusted score thresholds for each model using a previously reported approach [21]. This approach draws on the distribution of HMM-protein bitscores, phylogenetic relationships among proteins, and the relationship between phylogeny and bitscore to account for partial or divergent sequences and functionally dissimilar but related enzyme forms. All manually-set HMM thresholds can be found in Table S4. Generally, a pathway was considered present in a genome if at least one marker gene was found, as implemented in [12]. To reduce the incidence of false positives for a subset of pathways (formate oxidation, aerobic CO oxidation, and the Wood Ljungdahl pathway), we set a more stringent threshold, requiring that at least 2, 3, and 2, respectively, of marker genes to be present. Form I and II RuBisCO (Calvin Cycle) were identified as described below.

We next examined the distribution of each pathway as a function of the relative coverage of genomes encoding it. Per-genome relative coverage values computed above were merged with pathway presence information and summed by sample for each pathway (Fig. 2a). To examine the same metric by phylogenetic group (Fig. S2) relative coverage values of genomes with a given pathway were aggregated by phylum-level designation, then averaged within groups of samples (interface, anoxic zone, etc). Phyla which did not attain ≥ 1% mean relative coverage in any sample group were removed. Relative coverage values for potential methane producers (taxa containing the *mcr* operon or taxonomically affiliated with known methanogenic lineages) were visualized on a per-genome basis (Fig. 2b).

### Analysis of RuBisCO diversity and abundance

To determine the diversity of RuBisCO genes across the water column and sediments, all metagenomic assemblies were probed with a set of Hidden Markov Models (HMMs) describing the form I, II, II, II/II, and III RuBisCO respectively [59]. Stringent score cutoffs originally published with the models were applied and the best hit was selected for each individual protein. Scaffolds encoding above-threshold hits were gathered and clustered across samples using dRep at 95% ANI (*cluster -p 20 -pa 0*.*80 -sa 0*.*95*) [73]. A non-redundant, representative set of scaffolds encoding RuBisCO was created by selecting the set of cluster centroids that were binned (i.e., belonged to a draft genome) and/or were the longest. Reads from all samples were re-mapped to this unique set and mean coverage per sample was computed using CoverM, as described above. Only scaffolds with ≥50% breadth were considered ‘present’ in a given sample and retained for diversity analysis.

Next, putative taxonomy was assigned to all scaffolds encoding RuBisCO using a combination of bin taxonomy (for binned fragments), phylogenetic affiliation of co-encoded proteins, and manual curation. Sequence fragments clearly originating from eukaryotic organisms, such as algae and diatoms, were removed. Classification of newly-recovered RuBisCO sequences was obtained by placing them within a reference RuBisCO large subunit protein phylogeny [19]. Finally, the relative coverage of each representative scaffold encoding RuBisCO within each sample was computed by dividing mean coverage by the total coverage across all scaffolds. Relative coverage was first summed by phylum-level taxonomy of the encoding scaffold and then averaged across samples within the same lake compartment.

### CPR gene content analyses

Gene content of CPR bacteria was analyzed using a two-step *de novo* protein clustering pipeline [22]. First, proteins were clustered into ‘subfamilies’ and overlapping subfamilies were merged (–coverage 0.75). For sequences in each protein cluster, the most common KEGG annotation was determined alongside the frequency of this annotation. To determine statistically enriched functions by lake compartment, we drew on a previously developed computational pipeline that tests the frequency of occurrence of a function/protein cluster within genomes from a given lake compartment versus those from outside of that compartment [38]. To determine the degree of enrichment, a ratio describing the frequency of occurrence within a given compartment relative to the frequency of occurrence outside that compartment was computed, and statistical significance assigned using the Fisher’s Exact Test (the fisher_exact function in scipy.stats, alternative=‘two-sided’). In cases where no genomes outside of a given ecosystem compartment encoded the function of interest, the function was labeled ‘exclusive’ and no ratio was computed. All comparisons were corrected for false discovery rate using the multipletests function in statsmodels.stats.multitest (“fdr_bh”). Only protein families with more than 5 member sequences were considered.

### Phage-related analyses

A preliminary set of putative viral scaffolds was constructed by subjecting all assemblies (including the post 0.2 µm “70S” sample) to viral prediction using VIBRANT [74] (default parameters) and VirSorter2 [75] (*--include-groups dsDNAphage,ssDNA --min-length 5000 --min-score 0*.*5*). All potential viral scaffolds ≥50 kb (including potential provirus scaffolds) were subsequently passed to CheckV [76] (*end_to_end* program) and results were used to further refine the set. Specifically, if a given scaffold was found to have either zero “host genes” (as determined by CheckV) and 5 or more “viral genes”, or, alternatively, ≥1 “host genes” but a greater number of “viral genes” and no “provirus” call, the scaffold was assigned likely virus status. If a given scaffold had 5 times or more the number of host genes as viral genes, and was not considered a provirus, it was discarded. If a given scaffold had 5 times or more the number of host genes as viral genes, and was considered a provirus, it was assigned likely provirus status. If a given scaffold had fewer than 5 times the number of viral genes than host genes, or no viral genes but less than 4 host genes, it was flagged for manual curation regardless of provirus status. Manual curation was performed by examining taxonomic and functional affiliations of proteins using a combination of databases as well as BLAST.

To further refine the set of putative viral genomes, we programmatically scanned for evidence of eukaryotic viruses. First, all viral proteins were predicted using Prodigal (*-p meta*) [55] and compared against UniRef100 as described above for rps3-bearing scaffolds. Taxonomic information for best, above-threshold hits for each protein was collected, and those scaffolds with ≥25% of annotated proteins affiliated with eukaryotes or eukaryotic viruses were discarded. The remaining set of putative phage were clustered at 95% ANI using dRep (*-pa 0*.*80 -sa 0*.*95 --clusterAlg single*) [73] and cluster representatives were chosen on the basis of scaffold length. Phage scaffolds were considered circularized if marked 100% complete by CheckV (via ‘high confidence’ direct tandem repeats).

Putative phage scaffolds were integrated with those published previously from a subset of Lac Pavin metagenomes [26,27], many of which were manually curated to completion. Distribution of the integrated set of phages was determined by stringent re-mapping of sequencing reads to the combined scaffold set as described above for bacterial and archaeal genomes. Host predictions were made using previously reported methodology [71] that combines CRISPR-Cas targeting, taxonomic profiling of phage genes, and, for predicted prophages, taxonomic profiling of surrounding genomic regions. Where host assignment methods were conflicting, assignment based on CRISPR-Cas targeting was prioritized, followed by taxonomic affiliation of regions surrounding prophages, where relevant.

Finally, we re-predicted proteins for the refined set of phage genomes using Prodigal (*-m single*). To account for the possibility of alternatively-coded phages, predictions were made in genetic codes 11, 15, and 4, and coding density was compared. If an ‘alternate’ code (15 or 4) displayed an increase in coding density ≥10% relative to standard (11) code for a given phage [77], protein predictions in that code were retained. Otherwise, protein predictions in standard code were retained. Phage metabolic potential with depth was then analyzed as described above for CPR bacteria. Recovered phage *pmoC* genes were combined with those recovered from bacterial draft genomes, aligned using MAFFT and trimAl [78], and incorporated into a maximum-likelihood tree using IQTree. Predicted proteins were also used to compare newly-reported phages from Lac Pavin (viral_region_length ≥ 50 kbp, Table S9) to those from reference databases/previous publications [26,28,79] using vContact2 [80]. The network was created using default parameters with the ‘ProkaryoticViralRefSeq201-Merged’ reference database (*--pcs-mode MCL --vcs-mode ClusterONE*). Clusters reported by vContact2 were further classified as those containing only reference sequences, both Lac Pavin and reference sequences, and only Lac Pavin sequences. The network was visualized using Cytoscape v3.9.1 [81] in a prefuse force directed layout with a modified spring coefficient of 1×10^−6^. One large subnetwork containing multiple clusters was manually extracted from the broader network for the purposes of visualization.

## SUPPLEMENTARY FIGURES

**Figure S1.**
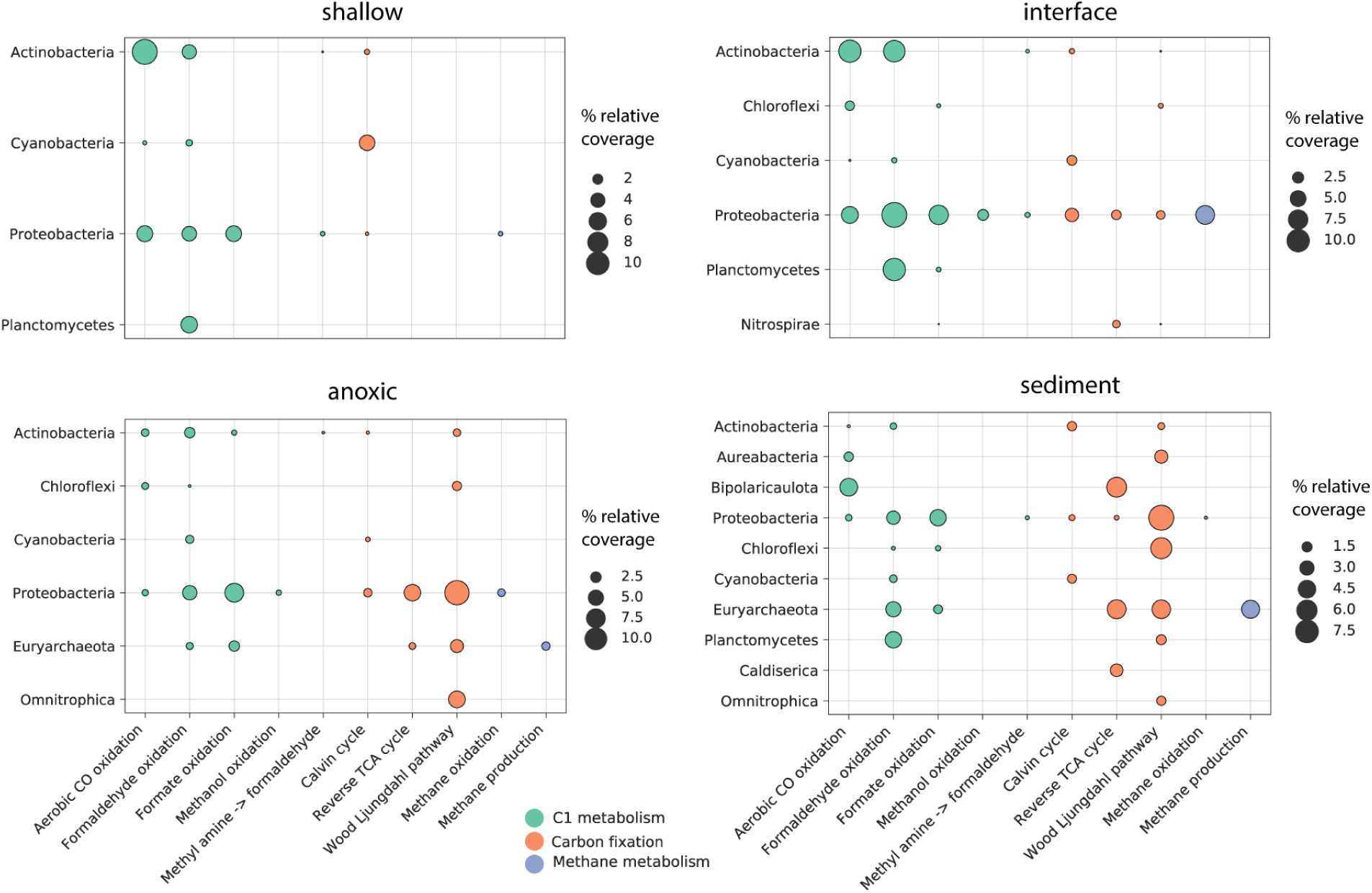
Genetic potential for select carbon metabolism across compartments Lac Pavin by phylogeny. **a)** Relative abundance of genomes encoding various pathways across metagenomic samples. Bubble size indicates the combined coverage of all organisms of a given phylum-level designation that encode a given pathway in each sample, normalized by the total coverage of all genomes in the quality-filtered, non-redundant set. This percent coverage is averaged across all samples within a lake compartment.

**Figure S2.**
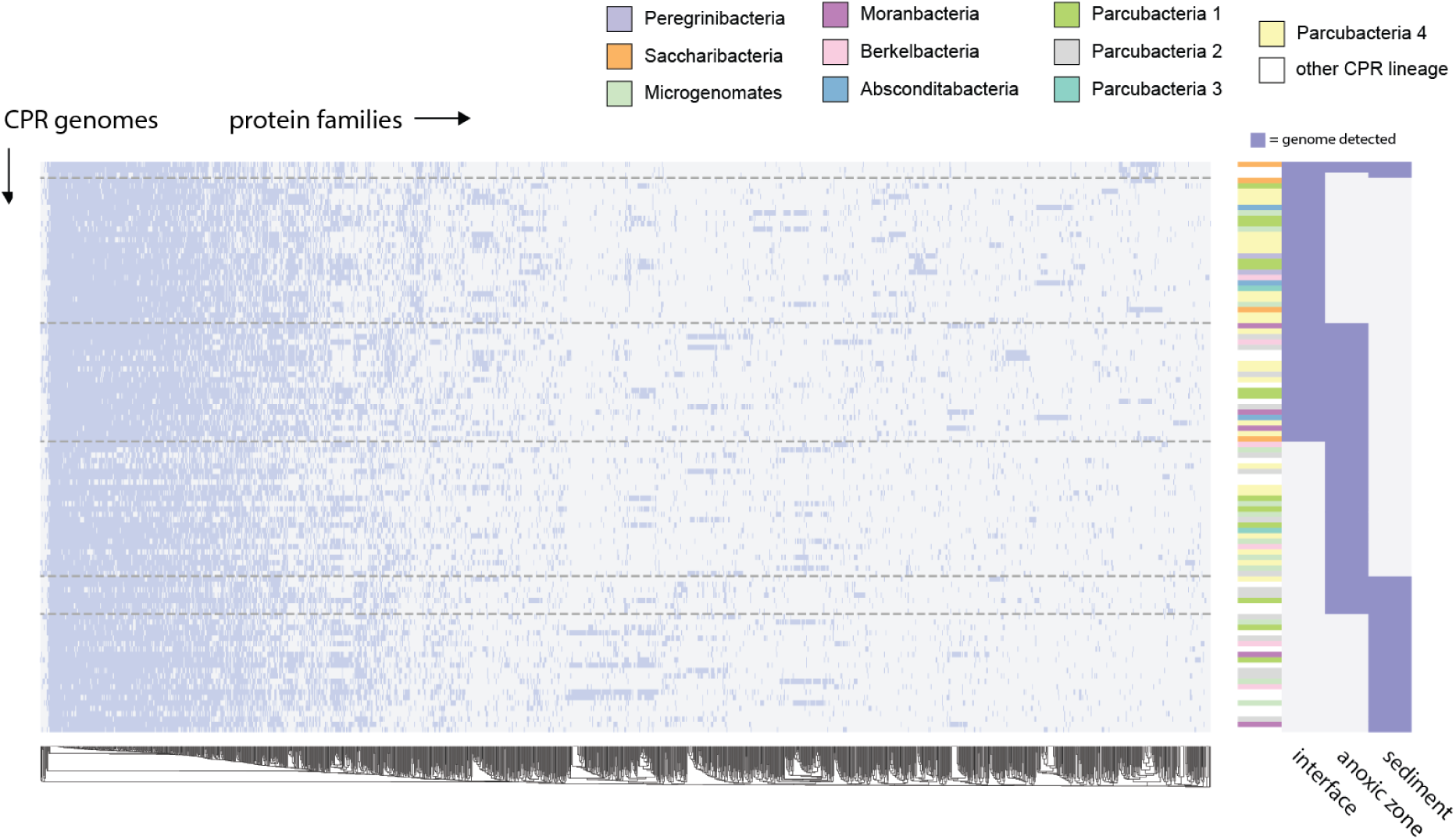
Proteome content in the CPR bacteria by depth **a)** Heatmap describing the presence/absence of ∼1,500 protein families (≥5 sequence members) across 106 genomes of CPR bacteria. Darker patches indicate presence, while lighter patches indicate absence. Protein families are hierarchically clustered by their distribution pattern across genomes. CPR genomes are organized by their distribution across the lake water column and sediments. In panel at right, darker patches signify that a given genome was present/detectable in a given compartment, whereas a lighter patch indicates absence/below detection threshold. Dashed lines distinguish sets of genome with similar distribution patterns, e.g. the bottom represents CPR bacteria detected only in the sediments.

